# The nucleosome acidic patch directly interacts with subunits of the Paf1 and FACT complexes and controls chromatin architecture *in vivo*

**DOI:** 10.1101/637223

**Authors:** Christine E. Cucinotta, A. Elizabeth Hildreth, Brendan M. McShane, Margaret K. Shirra, Karen M. Arndt

## Abstract

The nucleosome core regulates DNA-templated processes through the highly conserved nucleosome acidic patch. While structural and biochemical studies have shown that the acidic patch controls chromatin factor binding and activity, few studies have elucidated its functions *in vivo*. We employed site-specific crosslinking to identify proteins that directly bind the acidic patch in *Saccharomyces cerevisiae* and demonstrated crosslinking of histone H2A to Paf1 complex subunit Rtf1 and FACT subunit Spt16. Rtf1 bound to nucleosomes through its histone modification domain, supporting its role as a cofactor in H2B K123 ubiquitylation. An acidic patch mutant showed defects in nucleosome positioning and occupancy genome-wide. Our results provide new information on the chromatin engagement of two central players in transcription elongation and emphasize the importance of the nucleosome core as a hub for proteins that regulate chromatin during transcription.

## Introduction

The nucleosome, comprised of ~147bp of DNA wrapped around a histone octamer core, has long been regarded as a barrier to transcription in eukaryotes. However, evidence that the nucleosome serves as a recognition platform for factors that dynamically regulate chromatin structure has grown in recent years. While the amino-terminal tails of the four core histones H2A, H2B, H3 and H4 have been acknowledged as key regulatory components of the nucleosome, globular histone domains are also targeted by factors that alter chromatin structure and impact a diversity of nuclear processes (1). Notably, the nucleosome acidic patch, a negatively charged cavity shared by H2A and H2B, serves as an anchor point for a myriad of chromatin-binding factors. As demonstrated through crystallographic studies, proteins that bind to the acidic patch typically do so via a trio of arginines termed the “arginine anchor” (2). These proteins include the Latent Nuclear Antigen peptide (LANA) of Kaposi’s sarcoma virus, the H2A ubiquitylation module of Polycomb Repressive Complex 1, and the H2B deubiquitylation module of the SAGA coactivator complex (2–6). Through direct interactions, the acidic patch also regulates the binding and activities of ATP-dependent chromatin remodeling enzymes (7,8). Mutations that disrupt the acidic patch impair the binding and functions of proteins important for transcription (4,9), DNA damage repair (10,11), and chromosome segregation (12), and a recent search for mutations in gynecological carcinomas identified a missense mutation in the human gene *HIST1H2AB*, which leads to a substitution, H2A-E57Q, in a conserved acidic patch residue (13).

One crucial process regulated by the nucleosome acidic patch is the monoubiquitylation of a highly conserved lysine near the C-terminus of H2B (H2B K123 in *S. cerevisiae*, H2B K120 in *H. sapiens*) (5,9,14,15). In both yeast and humans, H2B ubiquitylation (H2Bub) is associated with transcription elongation and promotes Pol II processivity (16–18). H2Bub regulates chromatin structure, both directly and through its conserved role in promoting di- and tri-methylation of H3 at lysines 4 and 79 (17,19–23). At the level of nucleosomes, H2Bub has been shown to maintain nucleosome occupancy (19) and impact nucleosome positioning. For example, the chromatin remodeling enzyme Chd1 shifts hexasomes containing H2B K123ub more efficiently than those lacking the mark (24). H2Bub also plays a role in regulating higher order chromatin structure via inhibiting chromatin folding of nucleosome arrays (25).

H2B K123ub is catalyzed by the ubiquitin conjugase Rad6 and ligase Bre1 and is co-transcriptionally removed by the deubiquitylating enzymes Ubp8 and Ubp10 (26–32). In addition to these enzymes, the Polymerase Associated Factor 1 Complex (Paf1C) is required for H2Bub *in vivo* (33–35). In yeast, Paf1C contains five subunits, Paf1, Ctr9, Rtf1, Cdc73, and Leo1 (36–38). Through direct and indirect interactions with RNA polymerase II, Paf1C occupies the bodies of transcribed genes and regulates multiple co-transcriptional processes (39,40). The Rtf1 subunit of Paf1C plays a key role in activating H2B K123ub through its histone modification domain (HMD), which directly contacts Rad6 (33,34,41–43). In addition to Paf1C, the FACT histone chaperone complex has been implicated in regulating H2B K123ub as well as interacting with loci containing H2B K123ub (19,44). FACT is a conserved complex consisting of Spt16, Pob3, and Nhp6 in yeast and Spt16 and SSRP1 in humans (45). FACT disassembles and reassembles nucleosomes during transcription (46) and evicts nucleosomes from inducible promoters (47). FACT has multiple binding affinities for the nucleosome and has been shown to bind both histone dimers and tetramers (48–52). Recent studies have shown that Spt16 and Pob3 directly bind to H2A/H2B to disrupt nucleosome structure (53,54).

We previously demonstrated that the nucleosome acidic patch governs the H2B K123ub cascade, efficiency of transcription elongation and termination, and occupancy of Rtf1, Bre1, and Spt16 on chromatin (9). A recent study uncovered a direct interaction between the RING domain of Bre1 and the nucleosome acidic patch (15). Whether transcription elongation factors implicated in H2B K123ub directly engage the acidic patch or affect H2B K123ub through other mechanisms remained unclear. To address this question, we chose to apply an approach for investigating direct binding to the acidic patch in the context of cellular chromatin. With this objective, we site-specifically replaced amino acids in H2A with a photoactivatable, crosslinking-competent amino acid to identify proteins that directly contact the acidic patch *in vivo*. We detected crosslinking of H2A to Paf1C subunit Rtf1 and FACT subunit Spt16. Crosslinking of Rtf1 to H2A occurs independently of Bre1 and Rad6, suggesting that this interaction occurs independently of and possibly before catalysis. *In vivo* crosslinking experiments localized the H2A-interacting region of Rtf1 to the HMD, and *in vitro* binding assays revealed that the HMD binds to nucleosomes in a manner influenced by the nucleosome acidic patch. Consistent with the idea that the acidic patch serves as a hub for chromatin binding factors, MNase-seq analyses showed that the amino acid substitution H2A-E57A resulted in nucleosome occupancy and positioning defects genome-wide. Together, our data show that the nucleosome acidic patch is important for chromatin architecture and directly interacts with transcription elongation factors that regulate histone modifications.

## Materials and Methods

### Yeast strains, yeast growth media, and plasmid construction

The *S. cerevisiae* strains used in this study are listed in Table S1 and are isogenic to the strain FY2, which is a *GAL2*^+^ derivative of S288C (55). Yeast transformations were performed as previously described (56). For experiments involving BPA (p-benzoyl-L-phenylalanine), log phase yeast cultures were co-transformed with the tRNA/tRNA synthetase plasmid for BPA incorporation, pLH157/*LEU2* (43), and an ectopic H2A-expressing plasmid harboring an amber codon for BPA incorporation through nonsense suppression (57). *HTA1* was cloned into pRS426 as described (58). Gibson assembly and site-directed mutagenesis were used to introduce sequence encoding the HBH tag (59) prior to the *HTA1* stop codon to permit detection of full-length H2A proteins via a C-terminal tag. Site-directed mutagenesis of the parent H2A-HBH plasmid was used to introduce amber codons at positions 58 and 61 for BPA incorporation.

To create the strain inducibly expressing Rtf1-HMD_74-184_, oligonucleotides were used to amplify the coding region for amino acids 74 to 184 from Rtf1 and the fragment was subcloned into pAP39 (42) to generate pKB1463. Next, the segment expressing Myc-NLS-Rtf1-HMD_74-184_ was subcloned into pFA6a-KanMX (60), creating pKB1464, which adds a selectable marker for integrative transformation. Oligonucleotides with sequences flanking the genomic *RTF1* open reading frame were used to amplify the HMD region and the KanMX cassette from pKB1464, and this DNA was used to replace the native *RTF1* locus by integrative transformation. Inducible expression of the HMD was accomplished by replacing the native *RTF1* promoter with a galactose-inducible promoter from pFA6a-*TRP1*-*pGAL1* (60). Galactose-induced expression of the HMD promoted H2B K123ub (Figure S3). The final strain was co-transformed with the H2A-A61_BPA_-HBH plasmid and pLH157/LEU2 as above. Oligonucleotide sequences and plasmids are described in Tables S2 and S3, respectively.

### Photocrosslinking

50 mL of cells were grown to log phase (between 0.5 and 1.0 OD_600_) in SC-leu-ura + 2% dextrose and 1 mM BPA. For each culture, two samples of 12.5 OD units were pelleted and separately resuspended in 1 mL ddH_2_O. One sample was placed in the center of a 50-mL Falcon tube lid 2 cm below a UVGL-55 lamp to irradiate the cells at 365 nm and the other sample was processed as the non-crosslinked sample. Cells were exposed to UV light for 10 min and then processed for western blot analysis. For testing crosslinking of H2A-A61_BPA_-HBH to the HMD *in vivo*, strains were grown to early log phase in medium containing 2% raffinose and 1mM BPA, and HMD expression was induced by adding 2% galactose for 2 hr before UV crosslinking.

### H2A-HBH nickel pulldowns for mass spectrometry

100 mL of cells (KY2808 transformed with plasmids for BPA incorporation and encoding the following H2A derivatives: H2A-HBH, H2A-A61_BPA_-HBH, and H2A) were grown to OD_600_ 1.0. Cells were pelleted and resuspended in 4mL of ddH_2_O. Cells were UV-irradiated 1mL at a time for 10 minutes each. Cells were pelleted together and lysed by hand vortexing seven times (30s vortex alternating with one minute rest on ice) with ~500 µl glass beads in 500 µl lysis buffer (50 mM Tris-HCl, pH 8, 300 mM NaCl, 20 mM imidazole, 0.1% NP-40, and 8 M urea). Lysates were separated from glass beads and then cleared by centrifugation at 15000 × g for 10 min at 4 °C. 50 µl of lysate was saved for the input sample. ~450 µl was incubated with 100 µL pre-equilibrated (in lysis buffer) magnetic nickel beads (Qiagen #36111) overnight at 4 °C. The beads were washed four times with wash buffer (50 mM Tris-HCl, pH 8, 300 mM NaCl, 25 mM NaCl, 25 mM imidazole, 0.1% NP-40, and 8 M urea). Proteins were eluted in 50 mM Tris-HCl, pH 8, 300 mM NaCl, 1 M imidazole, and 8 M urea. All buffers contained 1X HALT protease inhibitors (Thermo Fisher). Eluates were analyzed by mass spectrometry at the Indiana University School of Medicine Proteomics Core Facility. The BPA-crosslinked samples were digested with LysC/Trypsin and then analyzed on a 50cm reverse phase column on a Orbitrap Fusion Lumos mass spectrometer. Multiple peptide-spectrum matches (PSM) were identified for Spt16 within the H2A-A61_BPA_-HBH +UV dataset; no matches for Spt16 were observed in the H2A-HBH or untagged H2A datasets. The PSM level identifications have a false discovery rate of equal to or less than 1%. To validate the quality of the Spt16 identifications, manual inspection of the MS/MS fragment ions was performed, which revealed good sequence coverage in both ion series further supporting the identification of Spt16. Identification of other H2A interactors by this approach awaits validation. We note that, despite the use of a buffer containing 8 M urea, high levels of common contaminants, such as translation factors and metabolic enzymes, were detected in the experimental and control samples (*i.e.* untagged H2A and wild-type H2A-HBH samples), potentially obscuring less abundant H2A interactors. Nonetheless, the number of peptides detected for Spt16 (5 peptides) was similar to the number of peptides detected for H2B (8 peptides), which is in close proximity to H2A-A61 (6), and H4 (5 peptides), which has been shown to contact the acidic patch through its N-terminal tail (6,58).

### Western blot analysis

Yeast cells were lysed by bead beating in trichloroacetic acid (TCA), as described previously (61). To detect H2A-HBH derivatives, proteins were resolved in SDS-polyacrylamide Tris-glycine gels (15% polyacrylamide) and transferred to nitrocellulose membranes. To detect crosslinking between H2A-HBH derivatives and Rtf1 or Spt16, proteins were resolved for 4 hr at 100 V on 4% −8% polyacrylamide gradient Tris-acetate denaturing gels (Novex, Life Technologies) and transferred to PVDF membranes. Because in general, the crosslinked species represent a small fraction of the total protein in question, we have provided both “dark” and “light” exposures of the western blots. For the light exposures, images of the blots were taken after ~30 second exposures. For the dark exposures, the same western blots were exposed for ~20 minutes and images were taken at the end of the duration.

Membranes were incubated with primary antibodies: anti-HA (Roche, #11666606001, 1:1000 dilution), anti-6X His tag (Abcam, #ab18184, 1:1000 dilution), anti-Spt16 (generous gift from Dr. Tim Formosa, 1:1000 dilution), anti-H2B K120 (recognizes yeast H2B K123ub (9,62), Cell Signaling 5546, 1:1000 dilution), anti-H2B (Active Motif, 39237, 1:5,000 dilution), anti-Rtf1 ((63) 1:5000), anti-H3 ((64), 1:3000 dilution) and then with anti-mouse or anti-rabbit secondary antibodies (GE Healthcare 1:5,000 dilution in 5% dry milk and 1X TBST). Proteins were visualized using enhanced chemiluminescence substrate (Thermo) and a ChemiDoc XRS digital imaging station (BioRad).

### Nucleosome pulldown assay

Plasmid-transformed strains were grown to an OD_600_ of 0.5 in 100 ml SC-leu-ura medium containing 1 mM BPA. Cells were pelleted by centrifugation at 1700 × g for 2 min at 4 °C then flash frozen in liquid N_2_. Cell pellets were thawed and lysed by bead beating in 500 µl lysis buffer [40mM HEPES, pH 7.9, 200 mM NaCl, 20 mM imidazole, 0.005% Tween 20 (v/v), 10% glycerol (v/v), 1x HALT protease inhibitors (Thermo Scientific, #78430)]. Lysates were cleared by centrifugation at 14,000 × g for 10 min at 4 °C. An aliquot of the cleared lysate (60 µl) was saved as input and the remaining volume was added to 80µl of Ni-NTA beads (Qiagen 36111) pre-equilibrated in lysis buffer. Beads were incubated with rotation overnight at 4 °C. Beads were washed three times with 750 µl 40 mM HEPES, pH 7.9, 200 mM NaCl, 25 mM imidazole, 0.005% Tween 20 (v/v), and 10% glycerol (v/v). Proteins were then eluted in 50 µl of the same buffer except containing 500 mM imidazole and subjected to western blot analysis.

### *In vitro* ubiquitylation assay

10 µl reactions were set up in the presence of reaction buffer containing 50 mM Tris-HCl, pH 7.9, 5 mM MgCl_2_, 2 mM NaF, 0.4 mM DTT, 4 mM ATP. In the following order, these factors were added to the reaction: 2.5 µg of reconstituted *Xenopus laevis* nucleosomes (generous gift of Song Tan), 15 pmol of purified recombinant HMD_74-184_ (43), 1.4 µg His-pK-HA-ubiquitin (43), 50 ng FLAG-hE1 (generous gift of Jaehoon Kim), 100 ng FLAG-yBre1 (generous gift of Jaehoon Kim), and 100 ng yRad6 (43). Reactions were stopped by adding 1X SDS loading buffer, boiling for 3 min in a thermocycler and flash freezing in liquid N_2_.

### Nucleosome electrophoretic mobility shift assay

Binding reactions containing 3 nM wild-type or acidic patch mutant *X. laevis* nucleosomes were incubated for 30 min on ice in binding buffer (10) containing 50mM Tris-HCl, pH 7.5, 150 mM NaCl, 2 mM arginine, 0.01% CHAPS (v/v), 0.01% NP-40 (v/v), and 1 µM BSA. Wild-type and acidic patch mutant nucleosomes were purified by anion-exchange HPLC (generous gift of Song Tan) and migrate similarly in native gels, indicating that the mutant nucleosomes are stable and not grossly misfolded. Recombinant HMD_74-184_ protein (43) was added at final concentrations noted in the figure legend. Reactions were loaded on 4-12% Tris-glycine native PAGE gels (Thermo, #XV04125PK20) and run for 90 min at 100V. DNA was visualized by SYBR Green staining (Thermo #S7563) in TE (25 min) followed by de-staining in TE (5-10 min) and imaging on a ChemiDoc XRS+ instrument (BioRad). The fraction of nucleosomal DNA shifted was quantified using Image Lab 6.0.1 (BioRad) and plotted against HMD concentration in R (https://www.r-project.org/). Each data point represents an average of at least three replicates, of which the standard deviation was calculated. Non-linear least squares regression was used to calculate K_d_ values using R.

### Micrococcal nuclease (MNase) sequencing

Mononucleosomes were prepared essentially as described in (65). Briefly, biological duplicates of wild-type and H2A-E57A mutant cells were grown in appropriate SC medium to an OD_600_ of 0.8 and crosslinked with formaldehyde at a final concentration of 1%. 100 mL of cells were pelleted, resuspended in FA buffer (50 mM HEPES-KOH, pH 8.0, 150 mM NaCl, 2.0 mM EDTA, 1.0% Triton X-100, 0.1% sodium deoxycholate), and lysed by bead beating. Following centrifugation, the chromatin-containing pellet was resuspended in NP-S buffer (0.5 mM spermidine, 0.075% IGEPAL, 50 mM NaCl, 10 mM Tris-Cl, pH 7.5, 5 mM MgCl_2_, 1 mM CaCl_2_), and then subjected to digestion by MNase (Thermo Scientific 88216) (Figure S1A). Mononucleosomal DNA was purified from treatments with 2.5 (low), 20 (mid), and 40 (high) U MNase digestions using agarose gel electrophoresis and Freeze N’ Squeeze Columns (BioRad 7326166). A fixed amount of MNase-digested, gel purified *Kluyveromyces lactis* DNA was added to each sample for spike-in normalization to assess occupancy changes using a method described in (66). Sequencing libraries were prepared using the NEBNext Ultra II kit (NEB E7645) and NEBNext Ultra sequencing indexes (NEB E7335) according to manufacturer’s instructions. Libraries were quantified using Qubit and TapeStation and pooled for paired-end sequencing on an Illumina NextSeq 500 (UPMC Children’s Hospital, Health Sciences Sequencing Core). Sequencing reads were aligned with HISAT2 (67,68) to the sacCer3 reference genome (69) and low quality reads were filtered using the SAMtools suite (70). Reads from biological duplicate samples were merged for downstream analysis. Reproducibility of the MNase-seq data for the two biological replicates, both wild type and H2A-E57A, is demonstrated through heatmaps, biplots, and a Pearson correlation plot in Figures S1B-D. Nucleosome positioning was analyzed genome-wide relative to published +1 nucleosome dyad positions (71) using the MNase option in the deepTools2 suite (72). Counts represent tag centers (3 bp) of uniquely mapped fragments 135 – 160 bp in length, counted in 1 bp windows and averaged over 50 bp. Heatmaps and metagene plots were generated using deepTools2 (72). Genome browser images were generated in Integrative Genomics Viewer (IGV 2.4.13) (73,74). Data presented here are lowly digested samples (2.5 U) to preserve and most accurately represent steady-state chromatin architecture.

In the course of this analysis, a duplication of chromosome VIII was detected in the H2A-E57A MNase-seq datasets. Using whole genome sequencing, we independently verified the disomy of chrVIII in this strain and screened additional H2A-E57A strains, generated by plasmid shuffle, for aneuploidies. In this way, we identified a strain with normal chromosome composition and subjected it, along with the original H2A-E57A strain, to H3 ChIP-seq analysis. A comparision of the H3 ChIP-seq datasets between the disomic and non-disomic H2A-E57A strains confirmed that chromatin architecture phenotypes were the same. We are, therefore, confident in our MNase-seq datasets. We removed chromosome VIII reads from MNase-seq analysis of both wild-type and H2A-E57A datasets to reduce any artifacts related to overrepresentation of chromosome VIII nucleosomes in meta-analyses. We note that in performing the H3 ChIP-seq experiment on two biological replicates of the newly derived H2A-E57A strain (individual colonies from a strain confirmed to have normal chromosome content by whole genome sequencing), one of the replicates became disomic for chrVII during growth of the culture. This demonstrates that H2A-E57A mutants are prone to aneuploidy, presumably through their chromosome bi-orientation defect (12).

### ChIP-seq

35 OD_600_ U of log phase wild-type and H2A-E57A mutant cells were crosslinked and sonicated in biological duplicate using the protocol described in (75). Histones were immunoprecipated from 1 μg chromatin and 1 μL of anti-H3 (Abcam, 1791) conjugated to 20 μl protein G magnetic beads (Invitrogen, 10004D) per reaction. Libraries were generated using the Ovation Ultralow v2 kit (NuGEN, 0344) and subjected to 50-bp single-end sequencing on an Illumina HiSeq 2500 at the Fred Hutchinson Cancer Research Center genomics facility. We used bowtie2 to align raw reads to the sacCer3 reference genome (69). Reads were then filtered using SAMtools (70). Bigwig files of input normalized ChIP-seq data were generated from the filtered bam files using deepTools2 (72) and dividing the IP data by the input data. Matrices for metaplots were generated in deepTools2 using the annotation file from (76). The metaplot in the final figure (Figure 5G) is an average of two biological replicates after assessing correlation between the two replicates. To do this, fastq files of raw reads were merged and the above pipeline was repeated on the merged fastq files.

## Results

### *In vivo* site-specific crosslinking reveals that the nucleosome acidic patch is a highly occupied binding hub

The nucleosome acidic patch is an essential site for promoting histone modifications associated with transcription elongation (5,9,14,15). Our goal was to identify transcription-associated proteins that bind the nucleosome acidic patch. We were particularly interested in proteins that mediate H2B K123ub, as we and others identified a role for the acidic patch in promoting this mark (9,15). To identify acidic patch-interacting proteins, we employed site-specific crosslinking using a photoactivatable, unnatural amino acid, ρ-benzoyl-phenylalanine (BPA) incorporated into H2A at positions within the acidic patch (77). Others have successfully used this system to detect nucleosome-nucleosome interactions during mitosis (58). To perform the crosslinking experiments, a wild-type yeast strain was transformed with a plasmid expressing an engineered tRNA and tRNA synthetase, which are required for BPA incorporation through amber suppression (57), and a second plasmid, which expresses an ectopic copy of *HTA1* containing an amber codon at the position of interest. Two amber codon mutant plasmids were generated for incorporating BPA into H2A, yielding H2A-Y58_BPA_ or H2A-A61_BPA_ tagged at the C-terminus with a 6XHis-Biotin-6XHis (HBH) tag (59). Positions 58 and 61 were chosen for BPA incorporation (Figure 1A; orange and blue residues) because these sites are located near amino acids required for H2B K123ub *in vivo* (Figure 1A; red residues) (9). Yeast transformants were exposed to long-wave UV radiation to induce BPA-mediated crosslinks. Western blot analysis of whole cell extracts from cells expressing the H2A BPA derivatives revealed several UV-specific bands (Figure 1B), indicative of interactions between unknown proteins and H2A residues Y58_BPA_ and A61_BPA_. Based on the crystal structure of the nucleosome (Figure 1A), we performed western blot analysis with an H2B antibody and detected the H2A-H2B interaction in extracts from the A61_BPA_ derivative, which confirmed that the BPA crosslinking experiments could detect expected interactions (Figure 1C lane 5 and Figure 1B lane 5, 34kD band). To determine whether the HBH-tagged H2A and H2A-A61_BPA_ derivative could be incorporated into nucleosomes, we performed a pull-down assay using nickel beads followed by western blot analysis (Figure 1D). Both forms of H2A-HBH were able to interact with H3, consistent with the incorporation of the ectopically expressed H2A proteins into nucleosomes.

**Figure 1.**
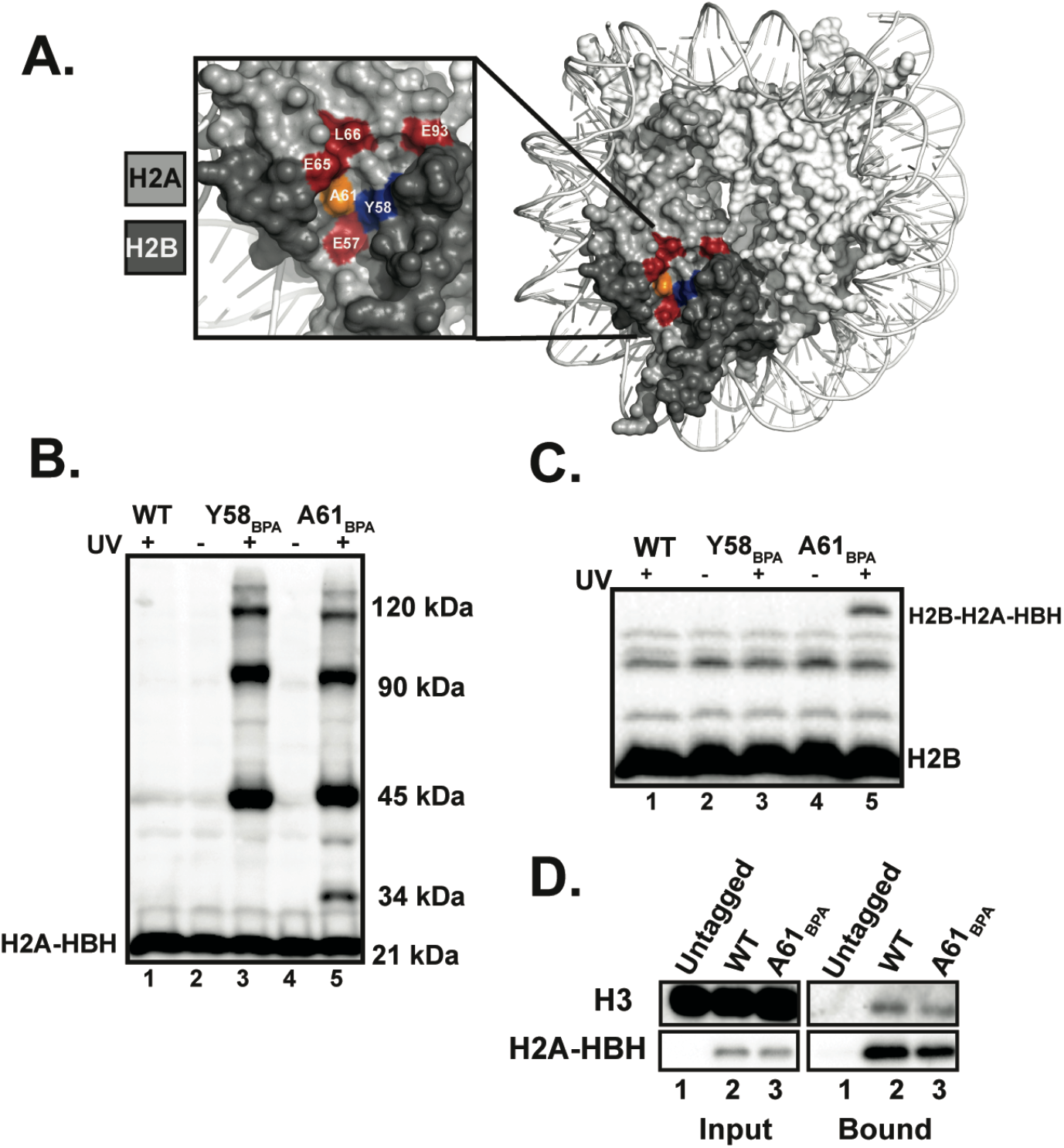
The nucleosome engineered for BPA crosslinking experiments. **A.** X-ray crystal structure of the nucleosome (PDB ID 1ID3 (6)) depicting sites of BPA incorporation in orange (A61) and blue (Y58). Residues in red are required for H2B K123ub *in vivo* (9). **B.** Representative western blot analysis of three biological replicates of HBH-tagged H2A probed with an anti-histidine tag antibody. Bottom band is the non-crosslinked H2A-HBH species and the upper UV-specific bands are crosslinked products of H2A-HBH. Cells ectopically expressed wild type (WT) H2A-HBH or the indicated BPA derivatives. Molecular masses are provided based on the broad range Biorad MW standard. **C.** Representative western blot analysis of three biological replicates of H2B in whole cell extracts prepared from cells exposed to UV light as indicated. Bottom band is non-crosslinked H2B. H2B crosslinked to H2A-HBH is indicated. **D.** Nickel pull-down, performed in biological duplicate, of H2A-HBH to assess interaction with H3 under non-crosslinking conditions. Western blots were probed with anti-his and anti-H3 antibodies.

### Paf1 complex subunit Rtf1 directly interacts with H2A *in vivo*

To determine the identity of H2A-crosslinked species, we performed western blot analysis of candidate proteins focusing on H2B K123ub-associated factors. We and others have demonstrated a requirement for an intact and accessible acidic patch for H2B K123ub *in vivo* and *in vitro* (9,15,78). Our previous work showed a marked decrease in the occupancy of Paf1C subunit Rtf1 at actively transcribed genes when residues within the acidic patch were substituted with alanine (9). To test if Rtf1 directly contacts the acidic patch *in vivo*, we performed western blot analysis of UV-treated cells expressing the H2A-BPA derivatives and a 3XHA-tagged form of Rtf1 (Figure 2A). Interestingly, we observed crosslinking of H2A to Rtf1 in a manner dependent on the location of the BPA residue within the acidic patch. While we reproducibly detected crosslinking of Rtf1 to the H2A-A61_BPA_ derivative, we did not observe crosslinking to the H2A-Y58_BPA_ derivative (Figure 2A; lanes 3 and 5). A61 is situated between two acidic residues, E57 and E65 (Figure 1A), and is perhaps more amenable to capturing certain interactions within the acidic patch compared to the Y58 location. We note that due to the inefficiency of amber suppression and the phenotypic consequences of reducing histone dosage, we chose to ectopically express the the H2A-Y58_BPA_ and H2A-A61_BPA_ derivatives over wild type H2A. This gives rise to cells with mixed pools of H2A and limits the levels of chromatin-associated, BPA-substituted H2A. As a result, the signals for the H2A-Rtf1 crosslinked species, and other H2A-crosslinked species, likely underestimate the interactions occurring within cells.

**Figure 2.**
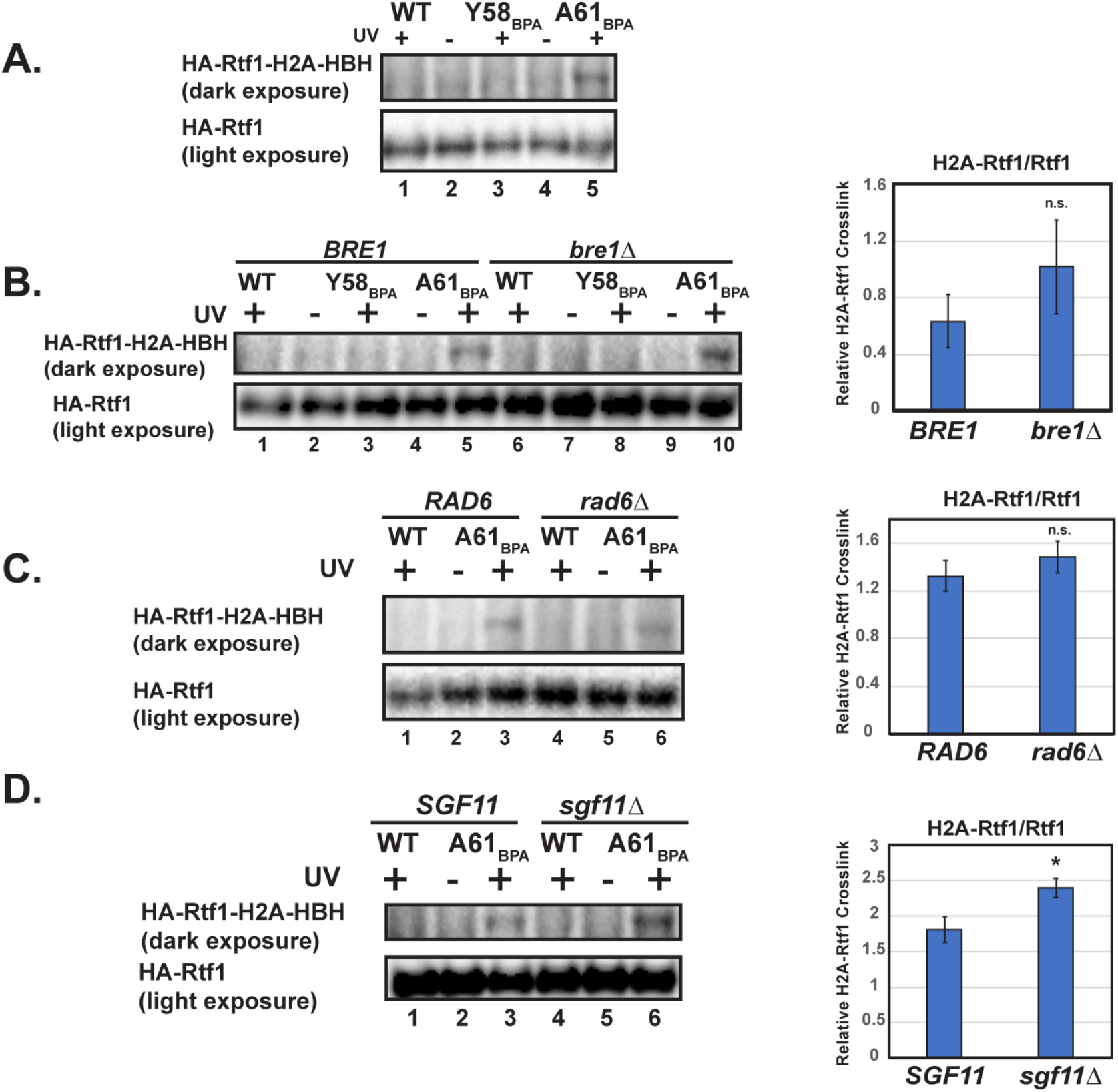
Rtf1 interacts with H2A independently of H2Bub enzymes. **A.** Western blot analysis of HA-Rtf1 in whole cell extracts prepared from cells expressing H2A-HBH (wild-type or BPA derivatives) and exposed to UV light where noted. Upper panel is a long exposure to detect crosslinked Rtf1 with H2A. Bottom panel is a shorter exposure of the same western blot showing total, non-crosslinked HA-Rtf1. **B-D**. Western analysis of H2A-HBH crosslinking to HA-Rtf1 in the absence of *BRE1* **(B)***, RAD6* **(C)** and *SGF11* **(D)**. Quantitation of crosslinked HA-Rtf1 normalized to total HA-Rtf1 levels is presented on the right. Error bars represent an SEM of three biological replicates and asterisks indicate a p value < 0.05 from a two-sample equal variance t-test. HA-Rtf1 was detected with anti-HA antibody.

### The Rtf1-H2A interaction is independent of the H2Bub enzymes

Due to the interdependent nature of chromatin association among Bre1, Rad6, and Rtf1 (18,43), we sought to determine whether the H2A-Rtf1 interaction required Rad6 or Bre1. Therefore, using strains deleted for *RAD6* or *BRE1*, we performed BPA-crosslinking experiments with the H2A-A61_BPA_ derivative and used the H2A-Y58_BPA_ derivative and wild-type H2A as controls. We found that the Rtf1-H2A interaction was still detectable in the absence of Bre1 or Rad6 (Figure 2B and 2C). The ability of Rtf1 to interact with H2A in the absence of Bre1 or Rad6 suggests that Rtf1 may interact with H2A before association of the enzymes required for catalysis.

A subunit of the H2B deubiquitylation module of the SAGA complex, Sgf11, has been shown to directly bind to the nucleosome acidic patch through X-ray crystallographic studies (5). Therefore, enzymes that place and remove the H2B K123ub mark recognize the same region of the nucleosome. To test if deletion of *SGF11* would influence the interaction between H2A and Rtf1, we performed crosslinking experiments in an *sgf11∆* strain. We observed a modest but reproducible increase in the H2A-Rtf1 crosslinked band in the absence of Sgf11 (Figure 2D). This result suggests that reducing competition to the acidic patch helps to increase binding of Rtf1 to H2A.

### The HMD directly interacts with nucleosomes *in vivo* and *in vitro*

Rtf1 is a multifunctional protein with several domains that have been previously characterized (Figure S2A) (39,41–43,62,79). The histone modification domain (HMD) is defined by regions 3 and 4 and is required for H2Bub and downstream modifications (41–43). Guided by this information, we sought to identify the specific regions within Rtf1 important for the H2A-Rtf1 interaction by performing BPA crosslinking experiments in yeast strains harboring integrated deletion alleles of *3XHA-RTF1*. Figure 3A and Figure S2 B-D show the different crosslinking species between Rtf1 mutant proteins and H2A detected by western blot analysis for the HA tag on Rtf1. We note that we observed several cross-reacting bands when analyzing the Rtf1 deletion mutants, some of which appear to crosslink upon UV-irradiation and some that do not. While we do not know the identity of these cross-reacting species, it is possible they represent modified forms of the Rtf1 mutant proteins. Focusing on the UV-dependent bands, the majority of *rtf1* deletion alleles did not disrupt crosslinking to H2A-A61_BPA_ (Figure S2 B-D). For example, we still detected crosslinking between Rtf1 and H2A when a domain within Rtf1 important for binding to Spt5 was mutated (39,41,79) (Figure S2A and S2C). This observation argues that even a transient association of Rtf1 with the Pol II machinery and chromatin can be captured by our crosslinking technique. In contrast, we found that the Rtf1∆4 protein, which lacks approximately the second half of the HMD, showed reduced crosslinking to H2A, whereas Rtf1∆3 protein, which lacks the first half of the HMD, crosslinked to H2A (Figure 3A). This observation indicates that a portion of the HMD is required for the H2A-Rtf1 interaction.

**Figure 3.**
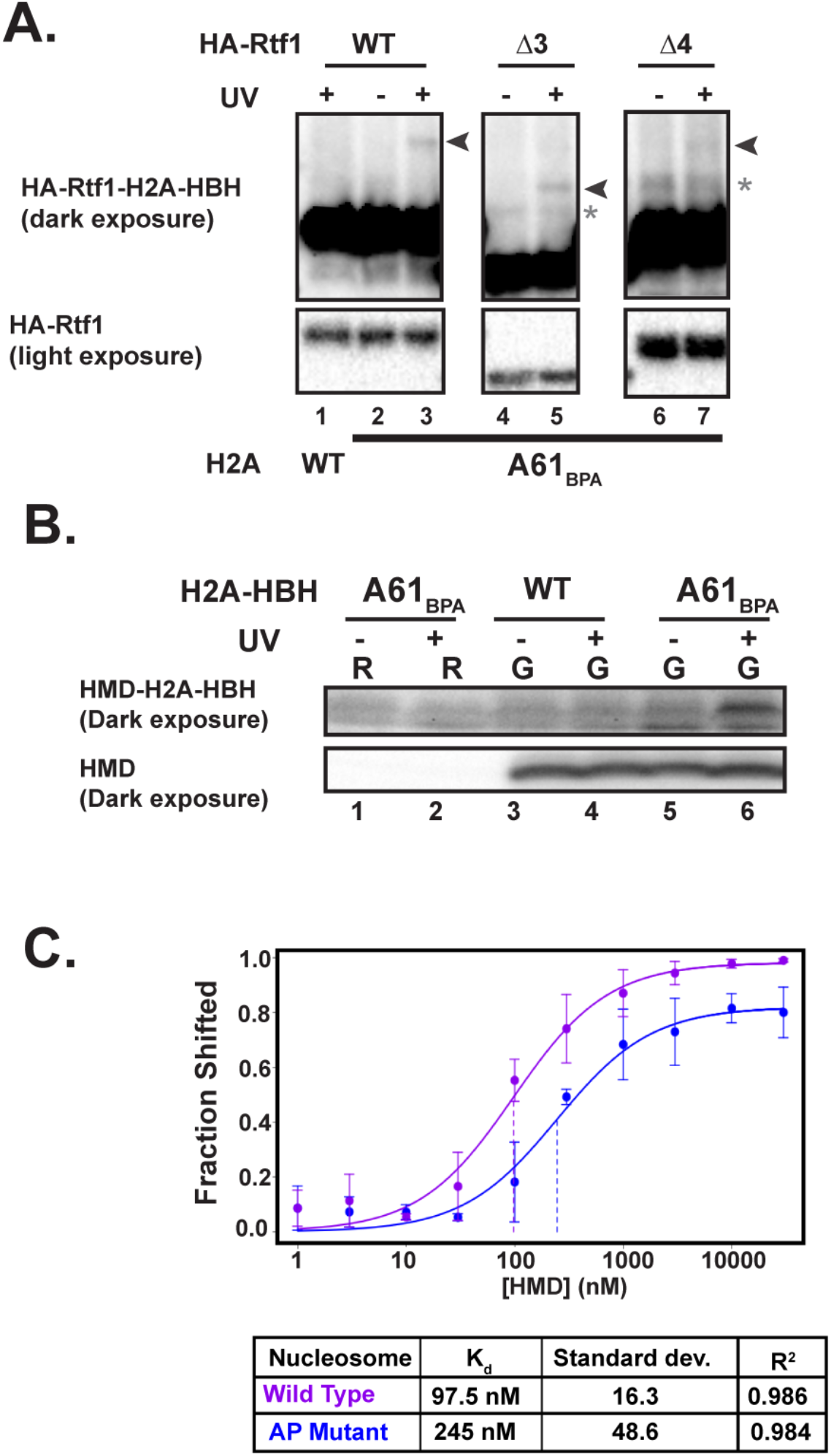
The Rtf1 histone modification domain binds the nucleosome acidic patch. **A.** Western blot analysis of the H2A-Rtf1 crosslink using an HA antibody to detect HA-tagged full-length Rtf1, Rtf1∆3, and Rtf1∆4. Arrows indicate crosslinked species. Asterisks indicate the presence of non-UV specific cross-reacting bands. Note that the Rtf1∆3 and ∆4 derivatives migrate faster than the full-length Rtf1. Upper panels are from a long exposure that shows both crosslinked and non-crosslinked HA-Rtf1. Bottom panels show a shorter exposure of the non-crosslinked HA-Rtf1 species only. All panels are from the same exposure (long or short) of the same western blot but are cropped for clarification purposes and all panels are representative westerns performed in biological triplicate. **B.** Western analysis of the HMD-H2A crosslink detected in extracts from cells grown in raffinose (R) or galactose (G) and in the presence or absence of UV exposure. Upper panel is a long exposure showing crosslinked HMD with H2A-HBH. Bottom panel is a short exposure showing non-crosslinked HMD. The blot was probed with an antibody against Rtf1, which recognizes the HMD. This was performed in biological duplicate. **C.** Quantitation of EMSA results showing binding of recombinant HMD to WT or acidic patch (AP) mutant *X. laevis* nucleosomes. The acidic patch mutant nucleosomes contain alanine substitutions at the following residues: H2A E61A, E64A, D90A, E92A. Each point represents an average of at least three replicates.

We previously showed that the HMD is sufficient to restore H2B K123ub to a strain lacking all five Paf1C subunits and argued that this small domain functions as a cofactor in the ubiquitylation process (43). Therefore, given its prominent role in promoting H2B K123ub, we tested whether the HMD alone could crosslink to the H2A-A61_BPA_ derivative. We generated a strain in which the endogenous *RTF1* gene was replaced with sequence encoding the HMD (Rtf1 residues 74-184) under the control of the *GAL1* promoter. In the presence of raffinose, the HMD was not expressed and H2B K123ub was not detected (Figure 3B bottom panel lanes 1-2 and Figure S3A). UV-exposure of raffinose-grown H2A-A61_BPA_ cells did not yield a crosslink-specific band that could be detected by the Rtf1 antibody (Figure 3B upper panel lanes 1-2). However, following 120 minutes of galactose induction, HMD expression and H2B K123ub were detected (Figure S3A), and importantly, a UV-specific and H2A-A61_BPA_-specific band was observed (Figure 3B upper panel lane 6). Detection of the HMD-H2A-A61_BPA_ crosslink (Figure 3B) provides independent validation of the Rtf1-H2A-A61_BPA_ crosslink (Figure 2) as these experiments involved different versions of Rtf1 (full length or HMD), different epitope tags (HA or myc) and different primary antibodies (anti-HA or anti-Rtf1). The formation of an HMD-H2A-A61_BPA_ crosslink indicates that the HMD makes direct contact with the nucleosome acidic patch *in vivo*.

As a complement to our *in vivo* studies, we asked if the HMD can directly bind to nucleosomes *in vitro* and if this binding requires an intact acidic patch. We performed electrophoretic mobility shift assays with recombinant wild-type or acidic patch mutant nucleosomes and increasing amounts of recombinant HMD. Interestingly, the HMD bound to the wild-type nucleosome with a Kd of ~100 nM (Figure 3C and Figure S3B). Upon incubation with the acidic patch mutant nucleosome, the binding efficiency was reduced nearly 2.5-fold to a Kd of ~250 nM (Figure 3C and Figure S3B). Together, these data show that not only does the HMD form a direct contact with Rad6 to promote H2B K123ub (43), but it also makes direct contact with the nucleosome acidic patch.

We previously showed the HMD can stimulate H2Bub in a reconstituted *in vitro* system containing purified *X. laevis* nucleosomes, human UBE1, yeast Rad6, yeast Bre1, and ubiquitin (43). Based on our nucleosome binding experiments, we attempted to ask if mutation of the acidic patch would block the stimulatory effect of the HMD in the reconstituted system. As we previously observed (43), addition of HMD_74-184_ to a reaction containing wild-type nucleosomes stimulated H2Bub (Figure S3C, lanes 2 and 3). In a reaction containing acidic patch mutant nucleosomes, the HMD was unable to increase the level of H2Bub. However, as previously reported (15), we found that the acidic patch mutant nucleosomes are strongly defective in supporting H2Bub even in the absence of the HMD with the final product level similar to that observed in a reaction lacking Bre1 (Figure S3C; compare lane 1 and lanes 4 and 5). Together with work from the Köhler lab (15), these results suggest that Bre1 requires an intact acidic patch to carry out H2Bub and this requirement prevents us from assessing stimulation by the HMD in the context of the acidic patch mutant nucleosome.

### BPA-crosslinking reveals an interaction between the acidic patch and FACT subunit Spt16

The common motif, as detected by X-ray crystallography, among proteins that bind to the nucleosome acidic patch is a group of three arginines that bind to the glutamic acids on the nucleosome, termed the “arginine anchor.” However, it has not yet been possible to predict proteins that bind the acidic patch bioinformatically (80). Thus, we sought to identify proteins that interact with the acidic patch using our BPA-crosslinking system followed by affinity purification under denaturing conditions and mass spectrometry.

We utilized the histidine tag to affinity purify HBH-tagged H2A proteins using nickel magnetic beads in the presence of 8 M urea (Figure S4A) and analyzed the eluates by mass spectrometry. Although these experiments were complicated by residual background signal that will require additional optimization to overcome (see Materials and Methods), we observed BPA-specific peptides for Spt16, a member of the FACT histone chaperone complex (Figure S4B). We obtained similar numbers of peptide hits for Spt16 (5 peptides) as for H2B (8 peptides) and H4 (5 peptides), both of which contact the acidic patch (see Figure 1C and (6,58)). The detection of peptides for Spt16 was intriguing to us because we previously reported that the acidic patch is required for proper histone occupancy as well as Spt16 occupancy on genes *in vivo* (9). A recent study also showed a reduction in co-immunoprecipitation between Spt16 and H2A from yeast cell extracts when certain acidic patch residues were substituted with alanine (81). We confirmed our mass spectrometry results by western blot analysis, which demonstrated UV- and BPA-specific crosslinking between Spt16-3XMyc and H2A-A61_BPA_ (Figure 4). Collectively, these data argue that the histone chaperone Spt16 directly contacts H2A within the acidic patch.

**Figure 4.**
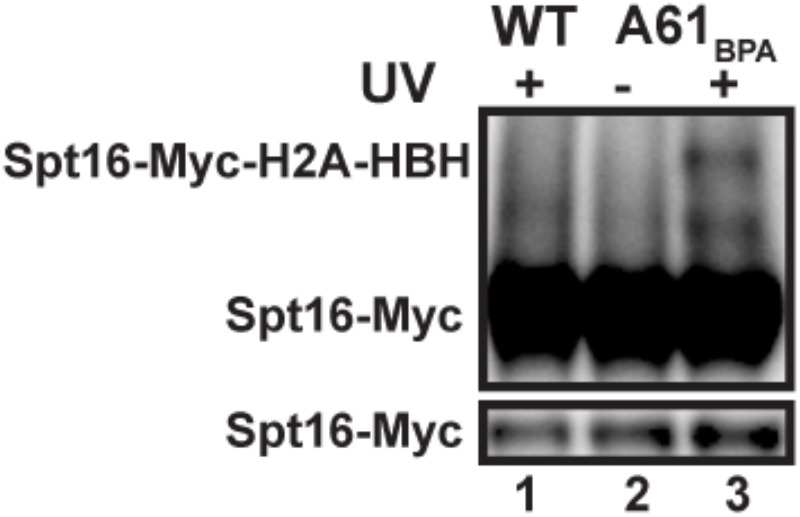
Spt16 interacts with the acidic patch. Representative western blot analysis of H2A-HBH crosslinking with Spt16-3XMyc probed with an antibody against the Myc tag performed in biological triplicate. Upper panel shows a long exposure and UV-specific crosslinking of Spt16-Myc and H2A-HBH. Bottom panel is a short exposure showing non-crosslinked Spt16-Myc. Both panels are from the same western blot.

Our genetic analysis revealed acidic patch residues, including H2A E57, important for Spt16 and Rtf1 occupancy on chromatin (9). To test if an intact acidic patch is required for crosslinking of the elongation factors to H2A, we created H2A-E57A, A61_BPA_ and H2A-Y58F, A61_BPA_ double mutants. While the H2A-E57A, A61_BPA_ double mutant was competent to crosslink to multiple proteins (Figure S5A), we noticed that it displayed a marked reduction in interaction with H3 as well as reduced UV-crosslinking with H2B (Figure S5B and S5C). The Y58F substitution caused a less severe defect. These observations are consistent with the idea that the acidic patch is important for proper nucleosome assembly or stability *in vivo*. Reduced crosslinking of Rtf1 to H2A-A61_BPA_ in the context of the E57A substitution further suggests that the interaction of Rtf1 with H2A may only occur when H2A is incorporated into nucleosome core particles (Figure S5D). In contrast to Rtf1 and H2B, the stronger crosslinking-dependent bands in Figure 1B and Figure S5A may correspond to proteins that interact with H2A outside the context of chromatin, as their levels do not decrease in the E57A background (Figure S5A). Additional studies will be needed to identify these proteins.

### The nucleosome acidic patch is required for proper nucleosome occupancy and positioning genome-wide

We have shown that the nucleosome acidic patch directly interacts with subunits of two transcription elongation factors, Paf1C and FACT, that regulate nucleosome occupancy and modification. Led by these findings, we set out to address whether mutations to the acidic patch alter nucleosome positions and occupancy genome-wide using MNase-seq analysis of a strain in which the only copy of H2A is the viable, H2B K123ub-defective E57A allele (9). The spike-in normalized MNase-seq data revealed a general reduction in nucleosome occupancy in the H2A-E57A mutant at all genes (Figure 5A-C) and a shift of nucleosomes toward the 5’ nucleosome-depleted region (NDR) in the H2A-E57A mutant (Figure 5B). Over gene bodies, this 5’-shift in nucleosome position in the mutant relative to wild-type increases from a modest 3 base pair shift, on average, at the +1 nucleosome to a more dramatic 12 base pair shift at the +4 nucleosome when all genes are considered (Figure 5B).

**Figure 5.**
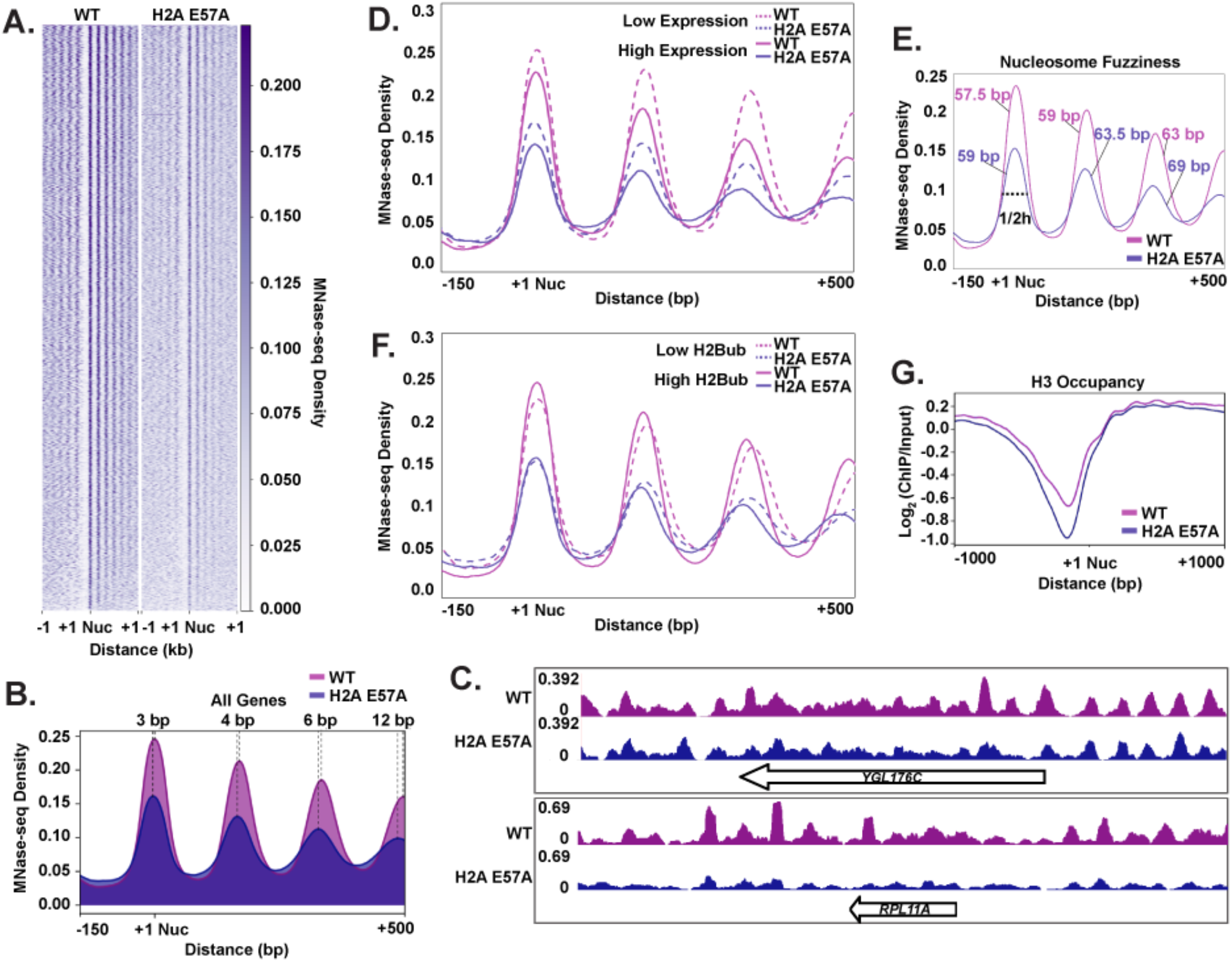
An intact nucleosome acidic patch is required for proper nucleosome occupancy and positioning. **A.** Heatmaps of nucleosome positions as determined by MNase-seq analysis of strains expressing plasmid-encoded wild-type H2A or H2A-E57A as the only source of H2A. Sequences are aligned to the +1 nucleosome of ORFs (71). Heatmaps represent averages of two biological replicates. **B.** Metaplot of nucleosome positions at all genes in the indicated strains. **C.** Spike-in normalized MNase-seq reads at two individual genes. **D.** Comparison of nucleosome positions in WT and H2A-E57A strains at lowly expressed and highly expressed genes (bottom 20% and top 20%) (82). **E.** Comparison of peak widths at ½ peak height in WT and the H2A-E57A mutant. Greater peak width suggests poorer nucleosome positioning on average across genes. **F.** Comparison of nucleosome positions at genes with high and low levels of H2Bub (bottom and top quintiles) (43). **G.** Metaplot of H3 ChIP-seq data at all genes. Experiments were performed in biological duplicate.

Due to the dynamic nature of chromatin during transcription, we asked whether this nucleosome shift in the H2A-E57A mutant was influenced by gene expression level. Therefore, we partitioned our MNase-seq data according to experimentally defined expression quintiles (82) and plotted read density averages in each of these subsets as metagene plots. In agreement with previous studies (83,84), nucleosome positions at the most highly expressed genes in the wild-type strain were shifted in the 5’ direction when compared to nucleosome positions at the most lowly expressed genes (Figure 5D, compare peak positions of the solid and dotted pink lines). The extent of nucleosome shifting in the H2A-E57A mutant was similar to that of wild-type when gene expression level was considered (Figure 5D, compare peak positions of the solid and dotted purple lines). In addition to this 5’ shift in position, we also observed a general loss of positioning at more 3’ nucleosomes in the mutant as evidenced by greater peak widths. For instance, at the +3 nucleosome (Figure 5E, compare pink lines to purple), the width at half peak height is 63 base pairs on average in the wild type compared to 69 base pairs in the H2A-E57A mutant. Increased peak width suggests “fuzziness” in positioning, or a pronounced inconsistency in nucleosome phasing within the population of mutant yeast compared to wild-type. While pervasive, the changes in nucleosome positioning, as measured by 5’ shift or peak width, observed in the H2A-E57A were not strongly correlated with gene expression level (Figure 5D).

H2B ubiquitylation levels generally correspond with transcription levels and nucleosome occupancy, and the acidic patch promotes H2B K123ub (9,15,19,85,86). Therefore, we asked whether changes in chromatin architecture relate to the reduced levels of ubiquitylation in the H2A-E57A mutant. We subset our MNase-seq data based on our published H2Bub ChIP-exo data (43) and found that nucleosome positions at highly ubiquitylated genes mirror the 5’ shift of highly expressed genes, consistent with the relationship between H2Bub and gene expression (Figure 5F) (17,35,87). We also observed, in agreement with previous data (19), that wild-type nucleosomes exhibit higher occupancy over highly ubiquitylated genes. This difference in occupancy is diminished in the H2A-E57A strain, likely as a result of the inability of this acidic patch mutant to promote placement of the H2Bub mark (Figure 5F).

Because non-histone proteins can block MNase digestion (88), we also analyzed chromatin structure in the H2A-E57A mutant by assessing genome-wide H3 localization using ChIP-seq (Figure 5G). Similar to the MNase-seq results, the mutant exhibited decreased H3 occupancy and shifted nucleosome positioning genome-wide. This analysis also revealed a widening of the NDR in the mutant strain. Together, these results demonstrate that the acidic patch influences nucleosome occupancy and positioning on active genes through mechanisms that likely depend on its multiple interactions with transcription elongation factors, histone modifiers, and other chromatin-associated factors.

## Discussion

In this report, we employed site-specific *in vivo* crosslinking to identify proteins that interact with the nucleosome acidic patch. Prompted by our discovery that the acidic patch is important for transcription-coupled histone modifications and transcription elongation (9), we focused our study on proteins involved in these processes. We identified direct interactions with the Paf1C subunit Rtf1 and the FACT subunit Spt16, adding to the compendium of proteins that influence the dynamic process of H2B ubiquitylation by contacting the acidic patch (Figure 6). *In vitro* nucleosome binding assays and *in vivo* BPA crosslinking experiments mapped the relevant nucleosome binding domain within Rtf1 to the HMD (Figure 3). Therefore, in addition to its interaction with Rad6 (43), the HMD, which can substitute for the complete Paf1C in stimulating H2B K123ub, is in direct contact with the nucleosomal substrate ultimately targeted by Rad6 and Bre1 for ubiquitylation. The crosslinking of Rtf1 to the acidic patch in the absence of *BRE1* and *RAD6 in vivo* (Figure 2) suggests that Rtf1 binds to the nucleosome independently of and possibly before engagement of these proteins. This ordering of events aligns with our recent ChIP-exo data (43), as well as earlier results (34,86), showing a partial loss of Bre1 and Rad6 occupancy in the absence of Rtf1. Our results support the hypothesis that a transcription elongation complex, Paf1C, can function as a cofactor in a histone modification process. Understanding how the HMD influences the binding of Bre1, which also contacts the acidic patch (15), will require additional studies.

**Figure 6.**
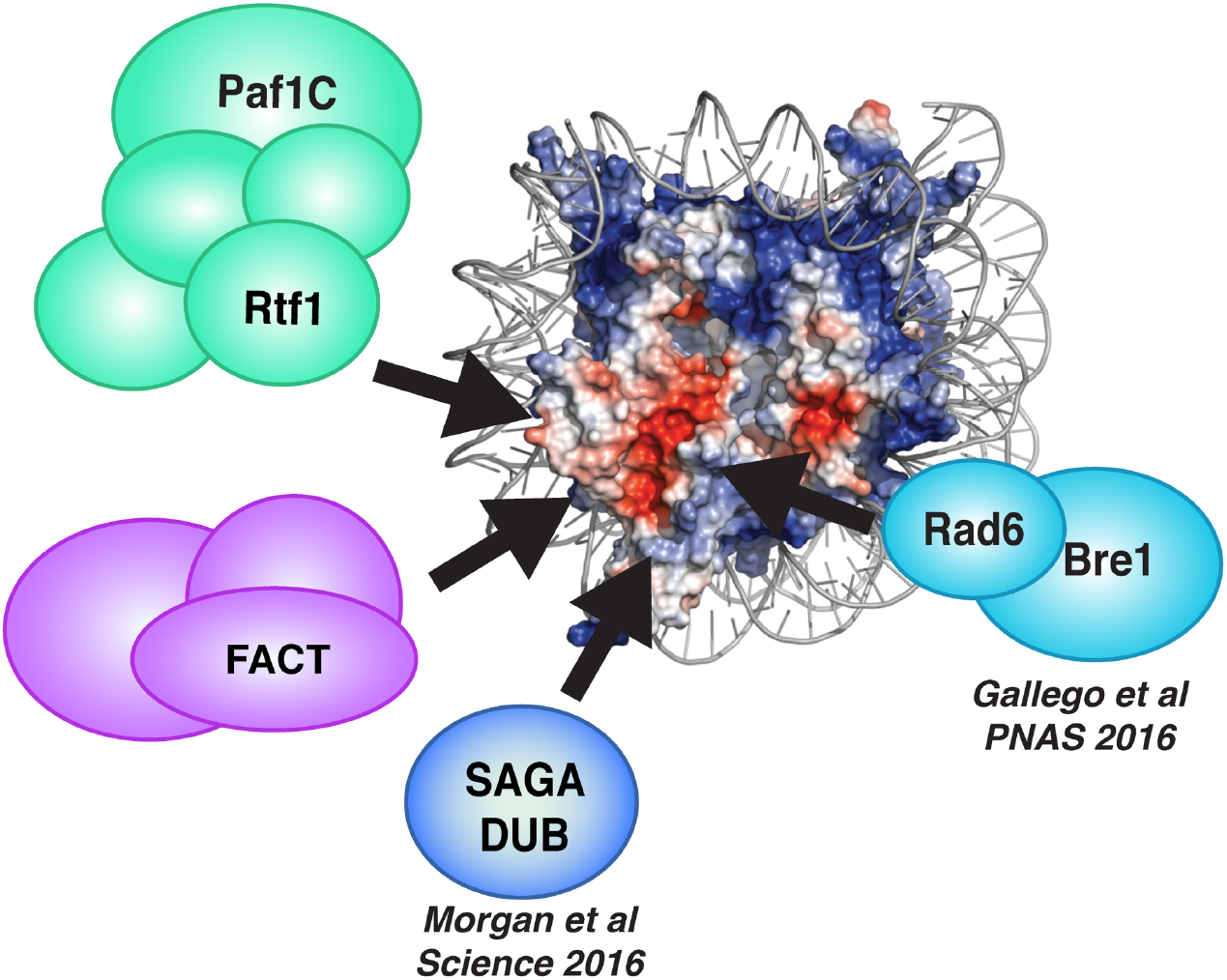
The nucleosome acidic patch is a key platform for regulating the chromatin landscape during transcription elongation. Electrostatic potential of the X-ray crystal structure of the nucleosome to highlight the nucleosome acidic patch (red). Proteins that bind the acidic patch and regulate H2B K123ub and/or transcription elongation, as revealed by this work or previous work, are indicated.

We previously showed that substitution of acidic patch residues causes reduced occupancy of both histones and Spt16 on active genes (9). Our BPA crosslinking analysis revealed that the nucleosome acidic patch residue A61 comes in contact with Spt16 (Figure 4). The histone occupancy defect in acidic patch mutants could be due, at least in part, to the reduction of Spt16 binding to H2A. Although no current crystal structures of Spt16 and histones show an interaction between Spt16 and the nucleosome acidic patch, it is likely that FACT recognition of the nucleosome is highly dynamic *in vivo* and interactions between Spt16 and the acidic patch may primarily occur in the context of active transcription. Indeed, a recent study found that transcription-disrupted nucleosomes enhance FACT binding to chromatin (89).

An important question is how protein binding to the acidic patch is regulated given the large number of proteins that interact with this surface of the nucleosome. When we deleted *SGF11*, which encodes a member of the SAGA DUB module that directly contacts the acidic patch (5), we observed an increase in H2A-Rtf1 crosslinking (Figure 2D), suggesting a competition between proteins that remove and stimulate H2Bub during transcription elongation. Given that Rtf1 and Spt16 crosslinking with H2A is sensitive to the location of the BPA residue, histones within the nucleosome may need to be properly positioned for these interactions to occur. Indeed, the intrinsic plasticity of the nucleosome is emerging as an important regulatory mechanism for chromatin transactions (90). For example, chromatin remodeling enzymes can transiently re-shape the nucleosome core via distortion during the remodeling reaction (91). It is thus possible that binding of FACT and Rtf1 at the acidic patch is regulated by changes to nucleosome structure that accompany transcription partly through the actions of chromatin remodeling factors and Pol II elongation complexes (92).

The nucleosome acidic patch is highly conserved across eukaryotes and several residues are essential for viability of yeast cells (93). In this study we assessed the impact of the viable H2A-E57A substitution on chromatin structure genome-wide using MNase-seq and uncovered defects in nucleosome positioning on gene bodies (Figure 5). We also found a widespread reduction in nucleosome and H3 occupancy in the acidic patch mutant. Because the acidic patch binds to many factors, the alterations in nucleosome occupancy and positioning detected in the H2A-E57A mutant are likely the net result of losing multiple interactions. However, we note that a genome-wide reduction in nucleosome occupancy has been reported for an *spt16* mutant and for an H2B mutant that lacks the site of ubiquitylation (H2B K123A) (19,94,95). It is also possible that the H2A-E57A substitution intrinsically renders the nucleosome unstable. Interestingly, two mutants defective in H2B K123ub, the H2B K123A mutant and a *rad6∆* mutant, exhibited a pattern of 5’ nucleosome shifting similar to what we observed in the H2A-E57A strain (19).

To date, most studies of factor binding to the nucleosome acidic patch have been performed *in vitro* under conditions that do not easily capture dynamic interactions, with the exception of H4-tail-acidic patch interactions during mitosis (58). Here, we present evidence of direct binding of two globally acting transcription elongation factors to the acidic patch *in vivo*. The recent discovery of many cancer-associated histone gene mutations (“oncohistones”) within the nucleosome acidic patch (13,96) further underscores the need to identify proteins that target the nucleosome acidic patch and mechanisms that regulate traffic to this critical interaction hub.

### Data Availability

All genomic sequencing data have been deposited in the GEO database under accession number GSE121543. Yeast strains and plasmids are available upon request.

## Supplementary Data

Supplementary Data have been submitted with the manuscript.

## Acknowledgments

We are grateful to Dr. Jaehoon Kim for providing purified hE1 and yBre1 for the *in vitro* ubiquitylation assay and to Dr. Song Tan for *X. laevis* wild-type and acidic patch mutant nucleosomes. We also thank Dr. Nathan Clark for providing the *Kluyveromyces lactis* type strain (NRRL Y-1140) as a spike-in control for genomics analyses. We thank Mitchell Ellison for assistance with bioinformatics analysis and Drs. Andrew VanDemark, Andrea Berman, Miler Lee, and Sarah Hainer, as well as members of their laboratories, for technical advice and support. We thank Dr. Toshi Tsukiyama for advice and support. We acknowledge technical support from Dr. Amber Mosley, Dr. Jason True and the Indiana University School of Medicine Proteomics Core Facility for mass spectrometry analysis and the UPMC Children’s Hospital Health Sciences Sequencing Core and FHCRC genomics core facility for sequencing of MNase-seq and ChIP-seq samples, respectively.

