## Supplementary Information for "The nucleosome acidic patch directly interacts with subunits of the Paf1 and FACT complexes and controls chromatin architecture *in vivo*"

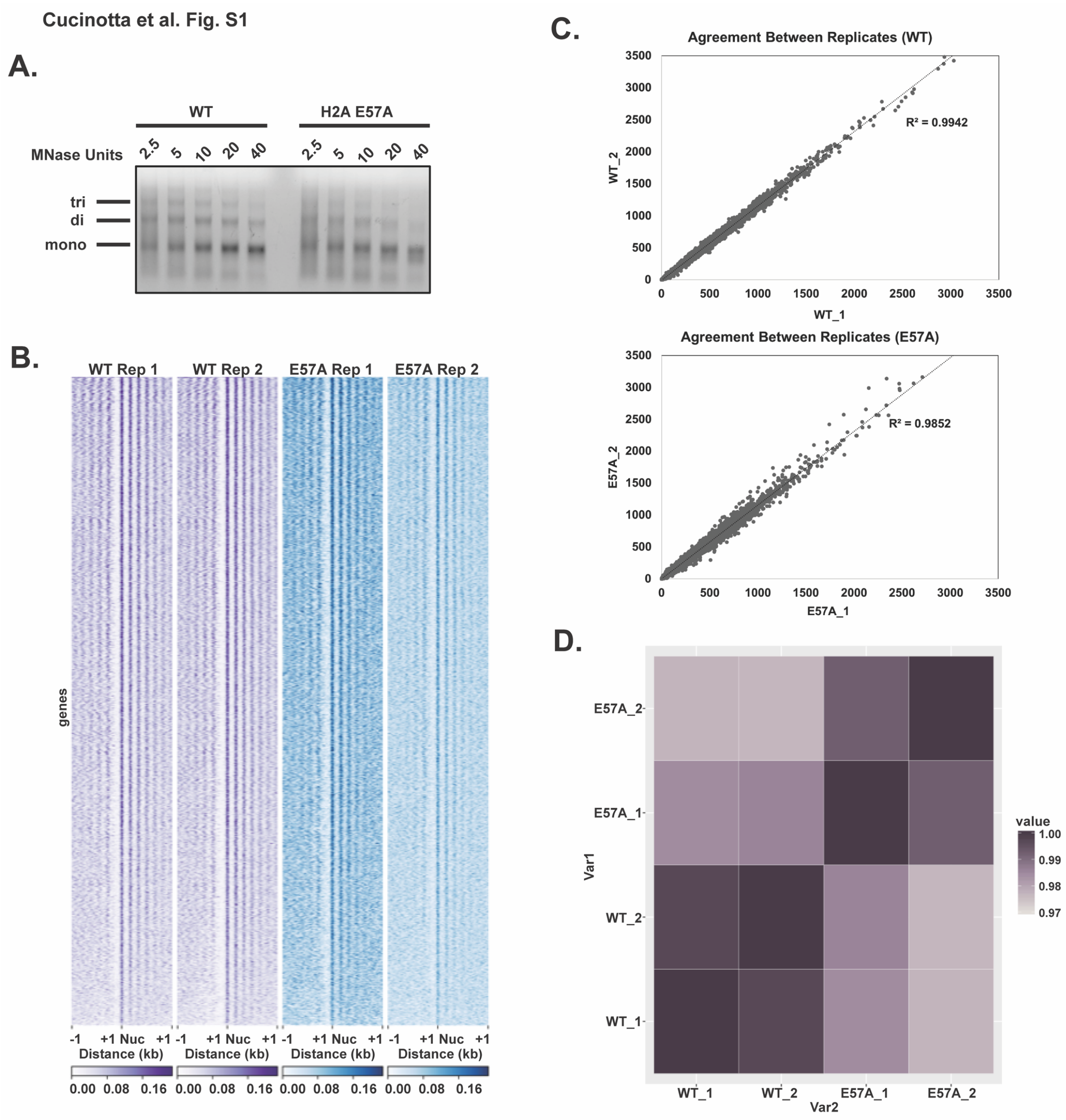


**Figure S1. MNase titration of cell lysates and analysis of MNase-seq data reproducibility**. **A.** Increasing units of MNase were added to cell lysates. Bottom band shows the mononucleosome species that was purified and subjected to library preparation and paired-end sequencing. **B.** Heatmaps of nucleosome positions determined in biological duplicate by MNase-seq analysis of strains expressing plasmid-encoded wild-type H2A or H2A-E57A as the only source of H2A. Sequences are aligned to the +1 nucleosome of annotated yeast genes. **C.** Biplots of MNase sequencing signal averaged over each annotated yeast gene comparing biological replicates of wild-type (top) or H2A E57A (bottom). $R^2$ values are indicated to the right of the line of best fit. **D.** Pearson correlation heat maps of MNase-seq data sets from all four samples. Darker color indicates higher correlation.

Cucinotta et al. Fig. S2

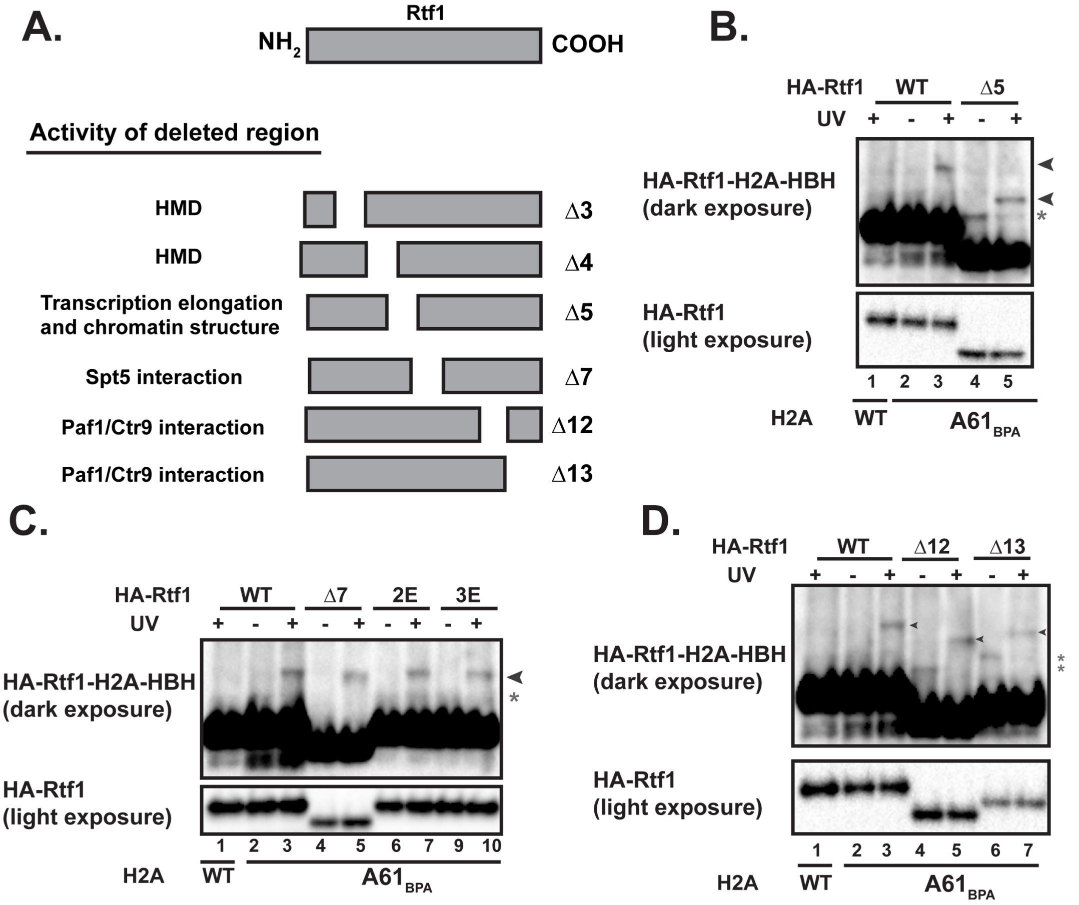


**Figure S2. Domain mapping of the H2A-Rtf1 interaction**

**A.** Diagram of Rtf1 deletion mutants and the function(s) of the corresponding domain that has been deleted (38). Western analysis of H2A-HBH crosslinking to wild-type HA-Rtf1 or HA-Rtf1 derivatives lacking region 5 **(B)**, lacking region 7 or containing glutamic acid substitutions for amino acids R273 and R288 (2E) or R251, R273, and K299 (3E) in region 7 **(C)**, or lacking regions 12 and 13 **(D)**. Arrows indicate crosslinked species. Experiments involving deletion mutants were performed in biological triplicate and experiments involving substitution mutants were performed in biological duplicate. Asterisks indicate the presence of non-UV specific cross-reacting bands. Note that the Rtf1 deletion derivatives migrate faster than the full-length Rtf1 as expected.

Cucinotta et al. Fig. S3

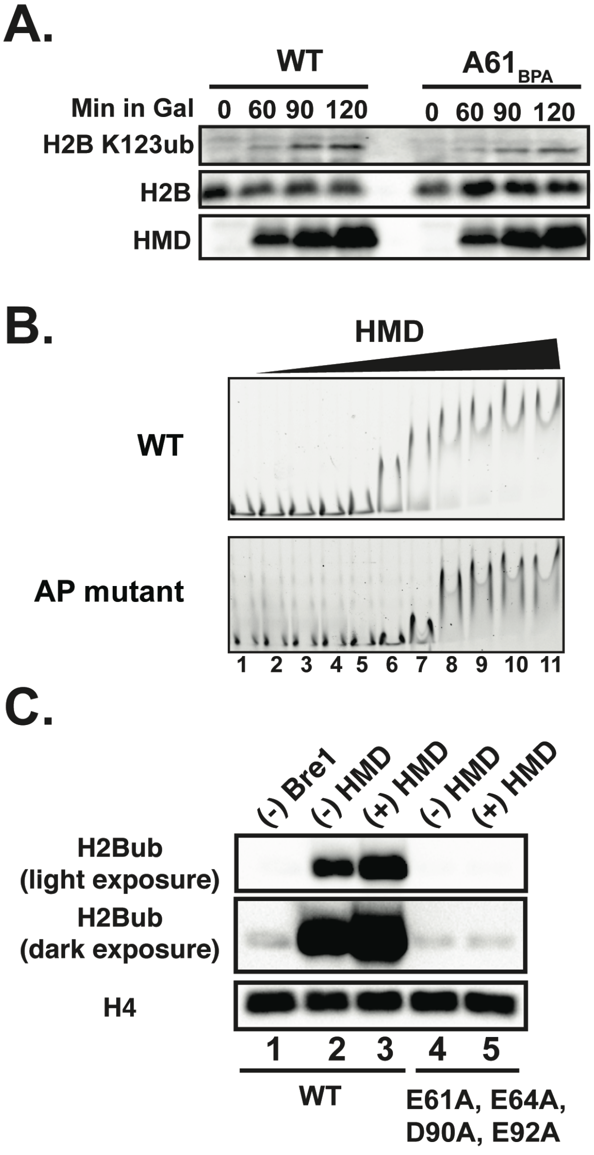


**Figure S3. Analysis of HMD binding to the acidic patch**

**A.** Time course of HMD induction in cells expressing the HMD under the control of the *GAL1* promoter. Western blots represent biological duplicate experiments, which were probed with antibodies against H2B K123ub, H2B, and Rtf1 (detects the HMD). **B**. EMSAs showing binding of purified, recombinant $HMD_{74\text{-}184}$ to wild-type and mutant recombinant *X. laevis* nucleosomes. Lane 1 has no HMD present. The following amounts of HMD were used in lanes 2-11: 1 nM, 3 nM, 10 nM, 30 nM, 100 nM, 300 nM, 1 μM, 3 μM, 10 μM, and 30 μM. **C.** *In vitro* ubiquitylation assay, performed in duplicate, of wild-type and acidic patch mutant nucleosomes. Except where indicated, all ubiquitylation reactions contained recombinant E1 (UBE1), E2 (Rad6), E3 (Bre1) and ubiquitin. Purified $HMD_{74\text{-}184}$ was added or omitted as indicated. The reaction in lane 1 lacked both Bre1 and HMD.

Cucinotta et al. Fig. S4

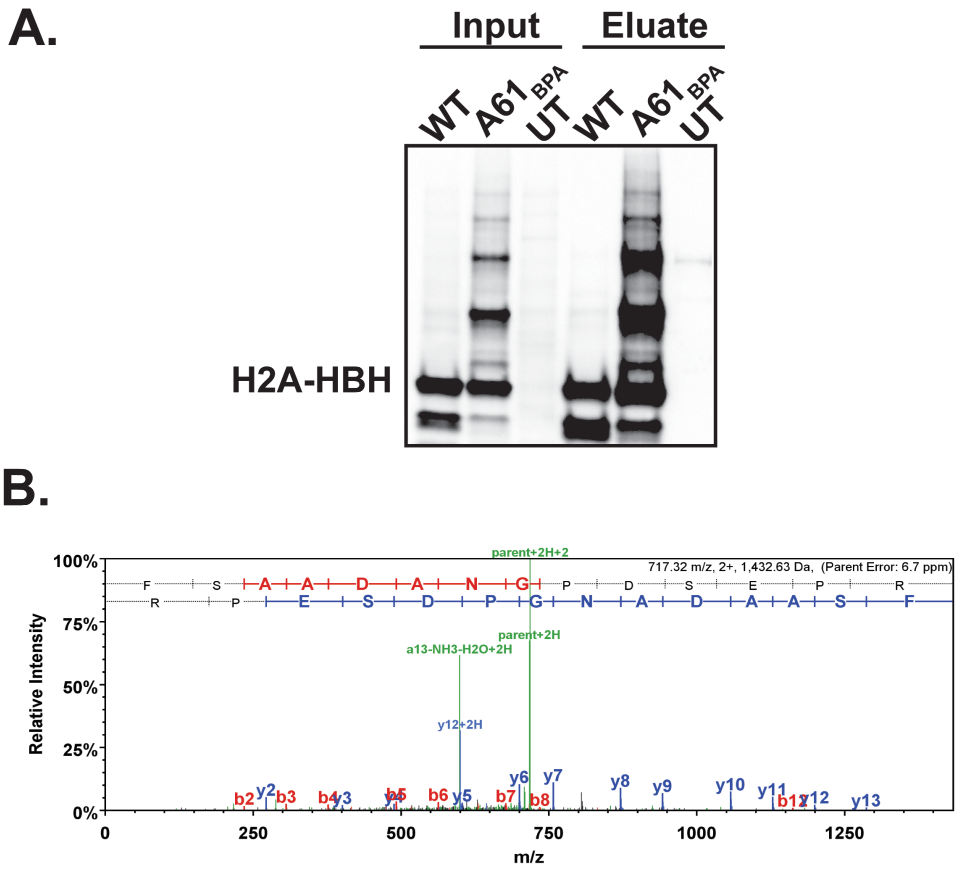


**Figure S4. Mass spectra of Spt16 peptides detected after pulldown of H2A-A61$_{BPA}$-HBH from UV-exposed cells**

**A.** Anti-his western blot analysis of eluates from nickel pulldown reactions performed in denaturing conditions for strains expressing H2A-HBH, H2A-A61$_{BPA}$-HBH, or an untagged H2A control (UT) and exposed to UV radiation. This experiment was performed in biological duplicate. **B**. Mass spectra of Spt16 peptides from one of the H2A-A61$_{BPA}$-HBH nickel pulldown experiments using extracts prepared from UV-treated cells under denaturing conditions.

Cucinotta et al. Fig. S5

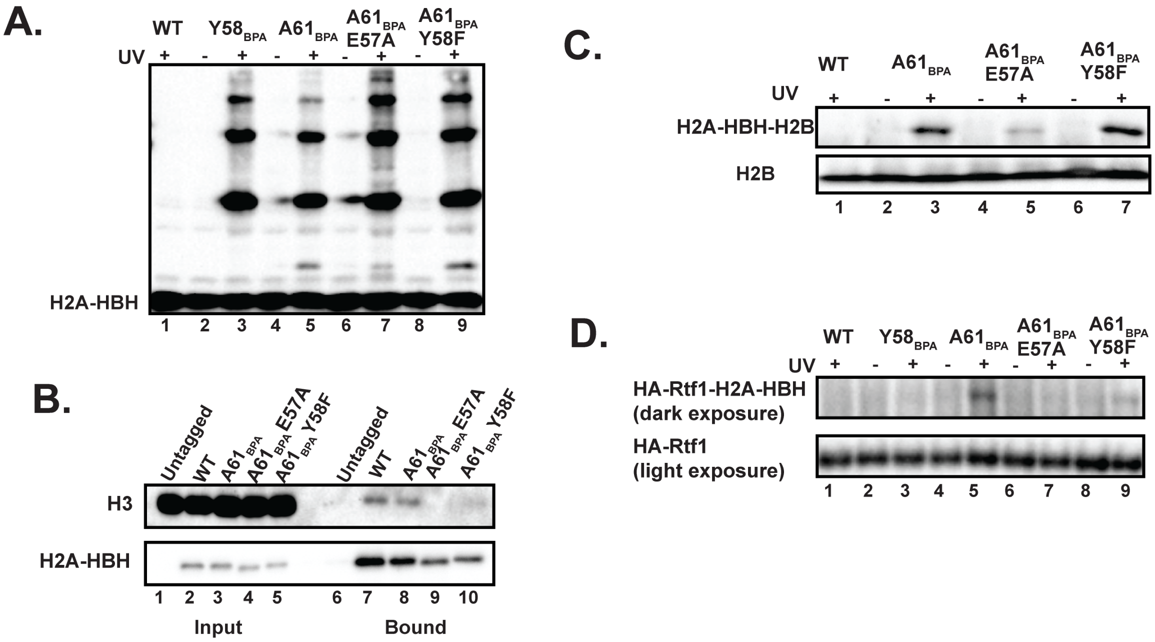


**Figure S5. An intact acidic patch is required for H2A-A61$_{BPA}$ to interact with H3**
**A.** Western blot analysis of extracts from UV-treated cells probed with anti-his antibody to detect the HBH tag appended to the indicated H2A derivatives. **B.** Western analysis of nickel pulldown experiments to analyze binding of the H2A-HBH derivatives to H3 under noncrosslinking conditions. Note that lanes 1-3 and 6-8 from these same blots are also shown in Figure 1D. **C.** Western blot analysis of H2A-H2B crosslinking from whole cell lysates. Top panel represents cross-linked species. Bottom panel is the non-crosslinked total H2B. **D.** Western blot analysis of H2A-HBH crosslinking with HA-Rtf1 in the context of H2A double mutants.

**Supplementary Table S1. Yeast Strains**

| Strain | *MAT* | Genotype |
|---|---|---|
| KY2808 | α | *his3Δ200 lys2-128δ leu2Δ1 ura3-52 3XHA-RTF1* |
| KY860 | a | *his3Δ200 lys2-128δ leu2Δ0 ura3Δ0* |
| KY2788 | α | *his4-912δ lys2-128δ ura3-52 trp1Δ63 3XHA-RTF1 bre1Δ::kanmx* |
| KY3047 | a | *his3Δ200 lys2-128δ leu2Δ1 ura3-52 3XHA-RTF1 rad6Δ::kanmx* |
| KY680 | α | *his4-912δ lys2-173R2 leu2Δ1 ura3-52 trp1Δ63 3XHA-rtf1Δ1* |
| KY2032 | a | *his4-912δ lys2-128δ leu2Δ1 trp1Δ63 ura3-52 3XHA-rtf1Δ3* |
| KY2033 | a | *his4-912δ lys2-128δ leu2Δ1 trp1Δ63 ura3-52 3XHA-rtf1Δ4* |
| KY1155 | a | *his3Δ200 leu2Δ1 ura3-52 trp1Δ63 3XHA- rtf1Δ5* |
| KY1157 | a | *his3Δ200 leu2Δ1 ura3-52 trp1Δ63 3XHA- rtf1Δ7* |
| KY2423 | a | *his4-912δ lys2-128δ leu2Δ1 ura3-52 trp1Δ63 3XHA-rtf1-R251E-R273E-K299E* |
| KY2424 | a | *his4-912δ lys2-128δ leu2Δ1 ura3-52 trp1Δ63 3XHA-rtf1 -R273E-R288E* |
| KY1159 | a | *his3Δ200 leu2Δ1 ura3-52 trp1Δ63 3XHA- rtf1Δ*12 |
| KY1420 | α | *his3Δ200 leu2Δ1 ura3-52 trp1Δ63 3XHA- rtf1Δ13* |
| KY977 (FY2365) | a | *his3Δ200 lys2-128δ leu2Δ1 ura3-52 SPT16-3XMYC* |
| KY3046 | α | *his3Δ200 lys2-128δ leu2Δ1 ura3-52 3XHA-RTF1 sgf11Δ::KANMX* |
| KY943 (FY406) | a | *(hta1-htb1)Δ::LEU2(hta2-htb2)Δ::TRP1 his3Δ200 lys2-128δ leu2Δ1 ura3-52* [pSAB6 = *URA3/C/A/HTA1-HTB1*] |
| KY3139 | α | *his4-912∂ leu2Δ1 trp1Δ63 rtf1::TRP1-GAL1p-NLS-MYC-HMD74-184::KanMX* |

**Supplementary Table S2. Oligonucleotides**

| Primer | Dir. | Sequence 5' → 3' | Ref. |
|---|---|---|---|
| Amplify *HTA1* 450 bp up and down ATG | F<br>R | ATCAGAGCTCGCGCTGTTCCAAAATTTTCGCC<br>ATCACTCGAGGCGTATATATATATACAAATATGCG | (54) |
| Back bone for gibson assembly to add HBH tag | F<br>R | TAAGATCGGTTCTGGTATTTTAAAG<br>TAATTCTTGAGAAGCCTTGG | This study |
| Insert for Gibson assembly to add HBH tag to H2A | F<br>R | AAGGCTTCTCAAGAATTATTAATTAACAGGGGTTCACATC<br>TACCAGAACCGATCTTAAGATCTATATTACCCTGTTATCC | This study |
| Y58TAG SDM | F<br>R | ACTTGACTGCTGTCTTGGAATAGTTGGCCGCTGAAATT<br>TCTAAAATTTCAGCGGCCAACTATTCCAAGACAGCAGTC | This study |
| A61TAG SDM | F<br>R | CTTGGAATATTTGGCCTAGGAAATTTTAGAATTAGC<br>CAGCTAATTCTAAAATTTCCTAGGCCAAATATTCCAAGAC | This study |
| Amplify $HMD_{74\text{-}184}$ for pAP39 cloning | F<br>R | GCTATTCCATATGGAAGAAGAAGCTAATCCTTTTCCCTTG<br>GGGAATTCTCATCATTCATTATCGCTGTAGTGACGGTTTTTCC | This study |
| Amplify $HMD_{74\text{-}184}$ for pFA6a-KanMX cloning | F<br><br>R | TGTAATTGTATTGCACTAATTTGTTGAGAGCACTATAGAAATGC CAAAGAAGAAGAGAAAGGTTGG<br>GCATGGCCTTGTTCCTTGGCACGAAACCATTCACATCACGAAT TCGAGCTCGTTTAAAC | This study |
| Integrate *GAL1p* at HMD | F<br><br>R | TGTAATTGTATTGCACTAATTTGTTGAGAGCACTATAGAAGAAT TCGAGCTCGTTTAAAC<br>GACTAGTTTGTGGAATACCAACCTTTCTCTTCTTCTTTGGCATT TTGAGATCCGGGTTTT | This study |

**Supplementary Table S3. Plasmids**

| Plasmid | Purpose | Derivation and reference |
|---|---|---|
| pCEC09/KB1473 | Untagged H2A | This study; (54) |
| pCEC21/KB1474 | WT H2A-HBH | Gibson assembly of pCEC09 |
| pCEC28/KB1476 | H2A-$Y58_{BPA}$-HBH | Site-directed mutagenesis of pCEC09 |
| pCEC29/KB1475 | H2A-$A61_{BPA}$-HBH | Site-directed mutagenesis of pCEC09 |
| pCEC30/KB1477 | H2A-$A61_{BPA}$-E57A-HBH | Site-directed mutagenesis of pCEC29 |
| pCEC31/KB1478 | H2A-$A61_{BPA}$-Y58F-HBH | Site-directed mutagenesis of pCEC29 |
| pLH157/*LEU2* | tRNA/tRNA synthetase containing plasmid | (40) |
| pKB1463 | Myc-NLS-Rtf1-$HMD_{74-184}$ | This study |
| pKB1464 | Myc-NLS-Rtf1-$HMD_{74-184}$-KanMX | This study |
| pAY01 | Plasmid containing *HTA1* and *HTB1* | (11) |
| pCEC02 | Plasmid containing *hta1-E57A* and *HTB1* | (11) |
